# A novel system for online fertility monitoring based on milk progesterone

**DOI:** 10.1101/248971

**Authors:** Ines Adriaens, Wouter Saeys, Tjebbe Huybrechts, Chris Lamberigts, Liesbeth François, Katleen Geerinckx, Jo Leroy, Bart De Ketelaere, Ben Aernouts

## Abstract

Timely identification of a cow’s reproduction status is essential to minimize fertility-related losses on dairy farms. This includes optimal estrus detection, pregnancy diagnosis and the timely recognition of early embryonic death and ovarian problems. On farm milk progesterone (P4) analysis could indicate all of these fertility events simultaneously. However, milk P4 measurements are subject to a large variability both in terms of measurement errors and absolute values between cycles. In this study, an innovative monitoring system based on milk P4 using the principles of synergistic control is presented. Instead of using filtering techniques and fixed thresholds, the present system employs an individually updated online model to describe the P4 profile, combined with a statistical process control chart to identify the cow’s fertility status. The inputs for the latter are the residuals of the online model, corrected for the concentration-dependent variability which is typical for milk P4 measurements. The objective of this paper is to present the developed methodology and give an indication of its use on farm. To this end, the system was validated on the P4 profiles of 38 dairy cows. The positive predictive value for luteolysis followed by estrus was 100%, meaning that the monitoring system picked up all estrous periods identified by the experts. Pregnancy or embryonic mortality was characterized by the absence or detection of luteolysis following an insemination, respectively. For thirteen cows, no luteolysis was detected by the system within the 25-32 days after insemination, indicating pregnancy, which was confirmed later by rectal palpation. It was also shown that the system is able to cope with deviating P4 profiles having prolonged follicular or luteal phases, which may suggest the occurrence of cysts. Future research is recommended for optimizing sampling frequency, prediction of the optimal insemination window and the establishment of rules to detect problems based on deviating P4 patterns.

**INTERPRETIVE SUMMARY. Progestrone-based online fertility monitoring. Adriaens.:** Milk progesterone based monitoring systems can be valuable for indicating fertility events in dairy cows, as the evolution of the milk progesterone levels can be related to acyclicity, estrus, pregnancy, and ovarian problems. We developed a system to translate raw progesterone measurements into clear information for the farmer, based on a mathematical model and a statistical control chart. In this way, a detailed image of a cow’s reproduction status on-farm can be obtained and fertility-related losses minimized.

## INTRODUCTION

Correct identification of a cow’s fertility status is important to further optimize reproductive performance in dairy cattle and consequently enhance farm profitability (Inchaisri et al., 2010). In order to minimize fertility-related losses, it is essential to obtain a complete image of a cow’s reproduction status as soon and accurately as possible. This should include information on the onset of cyclicity and estrus, successful inseminations, pregnancy, embryonic loss and the occurrence of ovarian abnormalities causing fertility problems (Friggens and Chagunda, 2005; Walsh et al., 2011). Today, systems based on external estrous symptoms fail to detect silent heats, nor are they suitable to identify other fertility events such as pregnancy or acyclicity. This presses the need for a system which combines (silent) heat detection with the detection of pregnancy, embryonic loss and the presence of ovarian abnormalities (Friggens and Chagunda, 2005).

Milk progesterone **(P4)** is widely accepted as a useful parameter to obtain a complete and direct image of a cow’s reproduction status (Friggens and Chagunda, 2005; Martin et al., 2013), while an automated system for online measuring milk P4 is already commercially available (Mazeris, 2010). One of the most important challenges is the interpretation of the raw sensor data (Rutten et al., 2013). Milk P4 measurements are subject to a large variability, partly caused by the measurement technique and calibration method (Adriaens et al., 2017), the sampling technique or the fat content in the milk sample (Pennington et al., 1981; Friggens et al., 2008). Additionally, P4 profiles also vary both within and between cows, for example in absolute values, slopes and lengths and often show irregular patterns (Meier et al., 2009; Blavy et al., 2016; Bruinjé et al., 2017a). Nevertheless, for milk P4 to be useful as an indicator for fertility events, the raw P4 measurements should be converted into specific information, or even better, ‘actions’ for the farmer. In this context, it is shown that both the variability of the measurements and the variability between estrous cycles should be taken into account (Friggens and Chagunda, 2005; Friggens et al., 2008; von Leesen et al., 2013). In general, this means that the monitoring algorithm should meet following requirements: (1) it should be robust against outliers and different levels of measurement errors; (2) it should be able to discriminate between luteal and follicular concentrations to indicate actions such as ‘inseminate’, ‘embryonic loss’, ‘cyst possible’ in an individualized way and (3) it should be automated and implementable on-line.

The closest attempt to this so far was published in 2005 by Friggens and Chagunda, (2005), who developed a biological model to predict the reproductive status of dairy cows based on milk P4. Besides different cow-specific factors, they used the smoothed level of the milk P4 obtained with an extended Kalman filter to account for the variability in the measurements. Next, this level is monitored and when it underruns a fixed threshold, the cow is indicated ‘in estrus’ and can be inseminated. Although this model has proven to be useful (Friggens et al., 2008) and easy to interpret, it has two major disadvantages, also identified in Friggens et al., (2008) and Bruinjé et al., (2017b): (1) the use of a smoothed P4 level causes a lag in the detection moment, dependent on e.g. sampling frequency and rate of luteolysis; (2) using a fixed threshold on the (smoothed) P4 level provides no flexibility to cope with the variability in individual P4 levels between cycles. For example, when a measurement fails at the crucial point of luteolysis, the smoothed level is not updated, and the next measurement of low P4 will be marked by this filter as ‘very unlikely’. Accordingly, the smoothed P4 level will adapt just moderately. Only when the following measurement is also low, it will move towards follicular P4 concentrations and ultimately undercut the fixed threshold (Bruinjé et al., 2017b). Moreover, this depends on milking and sampling frequency, and may occur for example more than three milkings after the actual occurrence of luteolysis, resulting in a large delay on the predicted ideal insemination moment. Additionally, if the follicular P4 concentrations are close to the fixed threshold, the detection lag will be even larger and successful timing of insemination even more unlikely.

The objective of this study was to develop a new approach to monitor fertility based on milk P4 measurements and the concepts of synergistic control (Mertens et al., 2009; Huybrechts et al., 2014; Maselyne, 2016). It was hypothesized that a system which combines a mathematical model to describe the P4 dynamics (Adriaens et al., 2017) with an online statistical control chart on the residuals of this model will allow on-line identification of the fertility status of a dairy cow and indicate important fertility events such as estrus, pregnancy and ovarian problems. Moreover, the proposed approach would be independent of fixed thresholds and does not imply a lag in the description of events like luteolysis, which would make the system more consistent and less dependent on the sampling rate and measurement errors.

## MATERIALS AND METHODS

### Experimental data

For this study, both milk P4 data and additional information on reproduction status were collected from the eligible cows on an experimental dairy farm in Geel, Flanders. Two trials were set up in the spring of respectively 2016 and 2017. In both periods, cows were milked automatically with an automated milking system of Delaval (VMS, Delaval, Tumba, Sweden) and were fed a mixed ration of grass and corn silage, supplemented with concentrates provided in the milking robot and through concentrate feeders. Because the procedure of data collection differed between the 2 periods due to the availability of an on-line milk P4 analyzer (Herd Navigator, Lattec, Hiller□d, Sweden) in the second period, each trial is described separately, and the information on cows and P4 data per period is summarized in Table 1. Nevertheless, for both trials, the P4 analyses were performed on mixed milk samples automatically collected from each milking following the procedure of the dairy herd improvement (DHI) protocol (ICAR International committee for animal recording, 2014). In 2016, the samples were frozen immediately at the test farm (-18°C), whereas in 2017, the samples were collected 3 times a week and brought to the Milk Control Center (MCC-Vlaanderen, Lier). Here, 2 mL subsamples were frozen after analysis of the milk constituents. At the end of each study-period, the P4 concentration of the thawed whole milk samples was measured with competitive ELISA, using a commercial milk P4 kit (Ridgeway, Gloucester, UK). The intra-assay CV were 29.4% and 7.8% in 2016, and 22.9% and 12.5% in 2017, for respectively low and high controls. The inter-assay CV were 43.8% and 19.7% (2016) and 38.0% and 18.5% (2017) for low and high controls respectively. Although these CV are high, they allow for clear discrimination between follicular and luteal P4 concentrations. A preliminary trial comparing with fresh and frozen variants of the same milk samples pointed out that there was no difference in P4 concentrations (results not shown). For further information on the sample collection, the P4 analysis and its accuracies, the reader is referred to Adriaens et al., (2017). Besides the milk P4 content, additional data of the cows were collected to confirm the onset of estrus, ovarian problems or pregnancy. To identify estrus or ovarian problems, the ovaries were checked by an expert veterinarian using an ultrasound scanner (A6v, Sonoscape Medical Corp., Shenzhen, China) equipped with a L745V 46MM 7.5 MHz linear transducer. The frequency of ultrasound to confirm the presence of a preovulatory follicle, a *corpus luteum* **(CL)** or ovarian cysts depended on the sampling trial (2016 or 2017) and is explained in the following paragraphs. In addition, the uterus tonus was checked through palpation during rectal examination by the expert as described by Bonafos et al. (1995). A follicular cyst was defined as a follicular structure (cavity) larger than 20 mm, either or not fully round and which persisted for more than 5 days on the ovaries in the absence of a CL which was confirmed by the lack of serum (< 1 ng/mL) or milk P4 (< 5 ng/mL). Luteal cysts were large structures (> 30 mm) with thick walls and often a cavity, persisting on the ovaries for more than 5 days after the expected luteolysis, producing P4 and resulting on milk P4 being higher than 5 ng/mL for more than 23 days. External estrous symptoms were scored visually, in particular the occurrence of standing heat, which typically occurs 26.4±5.2 hours (mean±SD) before ovulation (Roelofs et al., 2005), and metestrous bleeding, which is indicative for the end of the estrous period. Pregnancy was confirmed 40-50 days after insemination by rectal palpation and ultrasonography (KX5200V scanner 6.5 MHz, Kai Xin, Jiangsu, China).

**Table 1.**
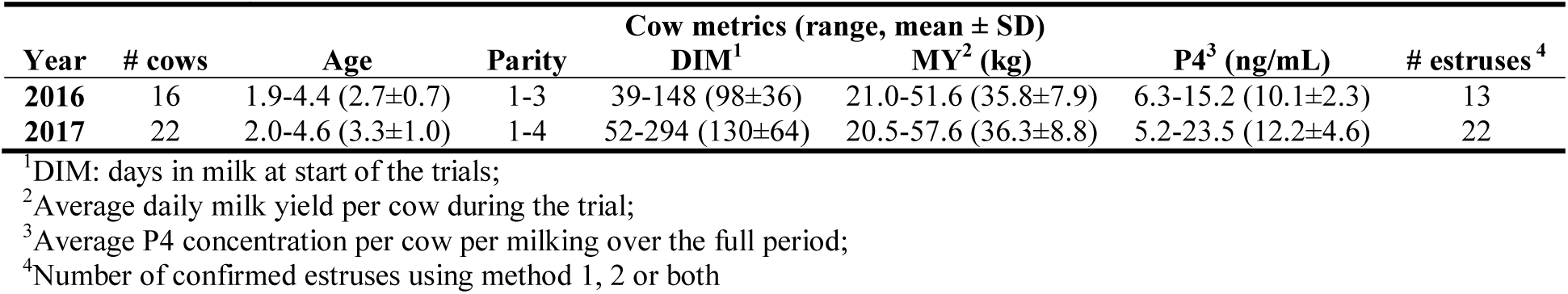
Summary of the cow metrics included in both studies

In 2016, the herd consisted of 52 lactating dairy cows. Two months before the trial, cows were selected based on their stage of lactation and reproduction status. Only animals not pregnant, cycling and beyond the voluntary waiting period of 30 days at the start of the trial were eligible, leaving 16 cows in the study. From these, the ovarian status was synchronized using the OvSynch protocol (Pursley et al., 1995). Sixteen days after the last GnRH injection, the ovaries of each cow were checked for the presence of a CL and luteal activity confirmed with a serum P4 sample > 1 ng/mL. Starting from day 19 after the last GnRH injection (typically 18 days from the previous induced ovulation), the uterus tonus and the growth of a preovulatory follicle were monitored daily until disappearance of the follicle. Seven to eleven days after the disappearance of the follicle, the presence of a CL was checked to confirm ovulation (Roelofs et al., 2010). If abnormal structures (e.g. thick walled structures or follicles > 20 mm) were detected on the ovaries, daily scanning continued until confirmation of cyst and treatment.

In 2017, the herd consisted of 58 cows. Two months before the start of the sampling period, the 28 cows which were beyond the voluntary waiting period and not confirmed pregnant were selected. The cows suspected to have cysts based on the online measured P4 profile were checked by an expert veterinarian and if needed, treated. At the start of the trial, cows confirmed pregnant or which didn’t (re)start normal cycling were excluded, leaving 22 cows. For these cows, the online raw milk P4 profile was assessed visually to detect the moment at which a consistent drop towards concentrations under 5 ng/mL started, staying below it for a period of at least 24 hours. This moment was named the Herd Navigator P4-drop **(HN-P4-drop)** and indicates the possible onset of estrus. Accordingly, synchronization of ovarian status was no longer needed. Between 15 to 50 hours after the HN-P4-drop, the ovaries were checked using ultrasonography for the presence of a preovulatory follicle. If the ultrasound image was not clear, scanning was continued daily upon detection of abnormalities or confirmation of ovulation. In the meantime, the uterus tonus was monitored. Between 7 to 11 days after the HN-P4-drop, the appearance of a CL confirmed ovulation.

Based on these data, confirmation of the onset of estrus was established in two different ways: (1) increased uterus tonus, showing a well contracted uterus upon rectal palpation (Bonafos et al., 1995) and absence of a CL, together with the presence of a preovulatory follicle of at least 13 mm diameter on one of the ovaries, which disappeared and was replaced by a CL on the same ovary 7 to 11 days later (method 1), in analogy with the method used by Michaelis et al., (2014); (2) the expression of standing heat (method 2) (Roelofs et al., 2010). All cows standing to be mounted were found to show metestrous bleeding 2 to 3 days later.

The available profiles were subdivided based on the presence of abnormalities detected by ultrasonography. Because in one case an abnormality occurred at the end of the trial, that profile was split up in two parts, either or not associated with the detected abnormality, and therefore included in the analysis twice. In this way, 33 normal profiles and 6 profiles from cows suffering from fertility issues were used to validate the developed system, originating from the 38 different cows. P4 data of 50.3±4.7 days per profile was included in the analysis. On farm, the system would start sampling in the *postpartum* phase, and the system is designed accordingly. However, because we sampled cows which were already cycling, none of the available profiles started in this first phase and the data of the incomplete cycle preceding the first follicular phase had to be discarded. As a result, P4 data from 3.5±3.9 days were excluded from the analysis. This is not a requisite for an online system, as on-farm monitoring can be initiated in the postpartum anestrus phase.

In total, 35 estruses were detected manually using the reference methodology also shown in Table 1. In 12 cases, both reference methods confirmed estrus. Twenty times only method 1 using ultrasonography counted (M1: uterus tonus, preovulatory follicle, CL), while the other 3 cases were observed by method 2 (M2: standing heat). Metestrous bleeding was noticed in 69% of the cases. From the 6 cows for which one or more fertility problems were detected, one cow first had a luteal cyst, which evaded spontaneously at the end of the trial. Another one was treated first for a follicular cyst and developed a luteal cyst hereafter. In 3 cows, a follicular cyst was identified and one animal displayed anestrus after showing a normal estrous cycle.

### Online model

In Adriaens et al. (2017), a model to characterize the P4 cycle was established, using a combination of two sigmoidal functions, together referred to as ‘full model’ (eq. 1).

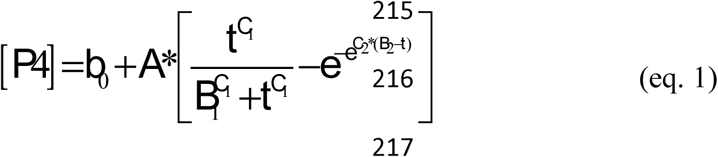

Each increasing part of a cycle, corresponding to the growth of a CL associated with an increase in P4, is described by a Hill function. The decreasing part associated with luteolysis can be characterized by a Gompertz function. *b*_*0*_ is the baseline P4 concentration when no CL is present. *A* represents the upper horizontal asymptote of the sigmoidal functions, and the *B* and *C* parameters define inflection points and slopes respectively. The parameter *t* represents the time within a cycle, starting from *t*_*0*_, i.e. parturition or the time of luteolysis of the previous cycle. In an on-line, on-farm system, this will be or the calving moment, or the moment a new cycle starts after luteolysis. For an elaborate description of the model and its parameters and how these functions were chosen, we refer to Adriaens et al. (2017). Besides the full model describing the P4 estrous cycle, the other stages of the reproduction cycles can also be identified using P4 and as such, this model can be incorporated in a system to monitor the fertility status online. A general overview of this online system is given in Figure 1. So, the inputs for the online monitoring system are the raw P4 values, from which four different stages can be deduced. Additional inputs are moment of parturition and inseminations.

(1) STAGE 1 = postpartum anestrus: After calving, P4 is produced only by the adrenal gland cortex, resulting in a low and constant basal milk P4 level, described as such by a constant (*b*_*0*_).
(2) STAGE 2 = luteal phase: With the onset of follicular activity, a dominant follicle will be selected and ovulates. The remaining theca and granulosa cells on the ovary form the *corpus hemorrhagicum*, which develops into an active CL producing P4. The corresponding increase in milk P4 from basal to luteal levels is described by an increasing Hill function. Each time a new measurement becomes available, the model parameters of that cycle (*b*_*0*_, *A*, *B*_*1*_, *C*_*1*_) are updated.
(3) STAGE 3 = luteolysis: When luteolysis occurs, the P4 production rapidly decreases back to basal levels. At this time, the second part of the model (i.e. the decreasing Gompertz function) is added to describe the P4 decrease and cycle characteristics can be calculated from the full model (*b*_*0*_, *A*, *B*_*1*_, *B*_*2*_, *C*_*1*_, *C*_*2*_) (= combination of increasing and decreasing part). After luteolysis, the P4 concentration is low (follicular phase) and the dominant follicle can develop further towards a preovulatory stadium and consequently ovulate. Steps 2 and 3 are repeated until a successful insemination establishes gestation
(4) STAGE 4 = pregnancy: In this case, the increase in P4 production is not followed by luteolysis and the P4 concentration remains high.

**Figure 1.**
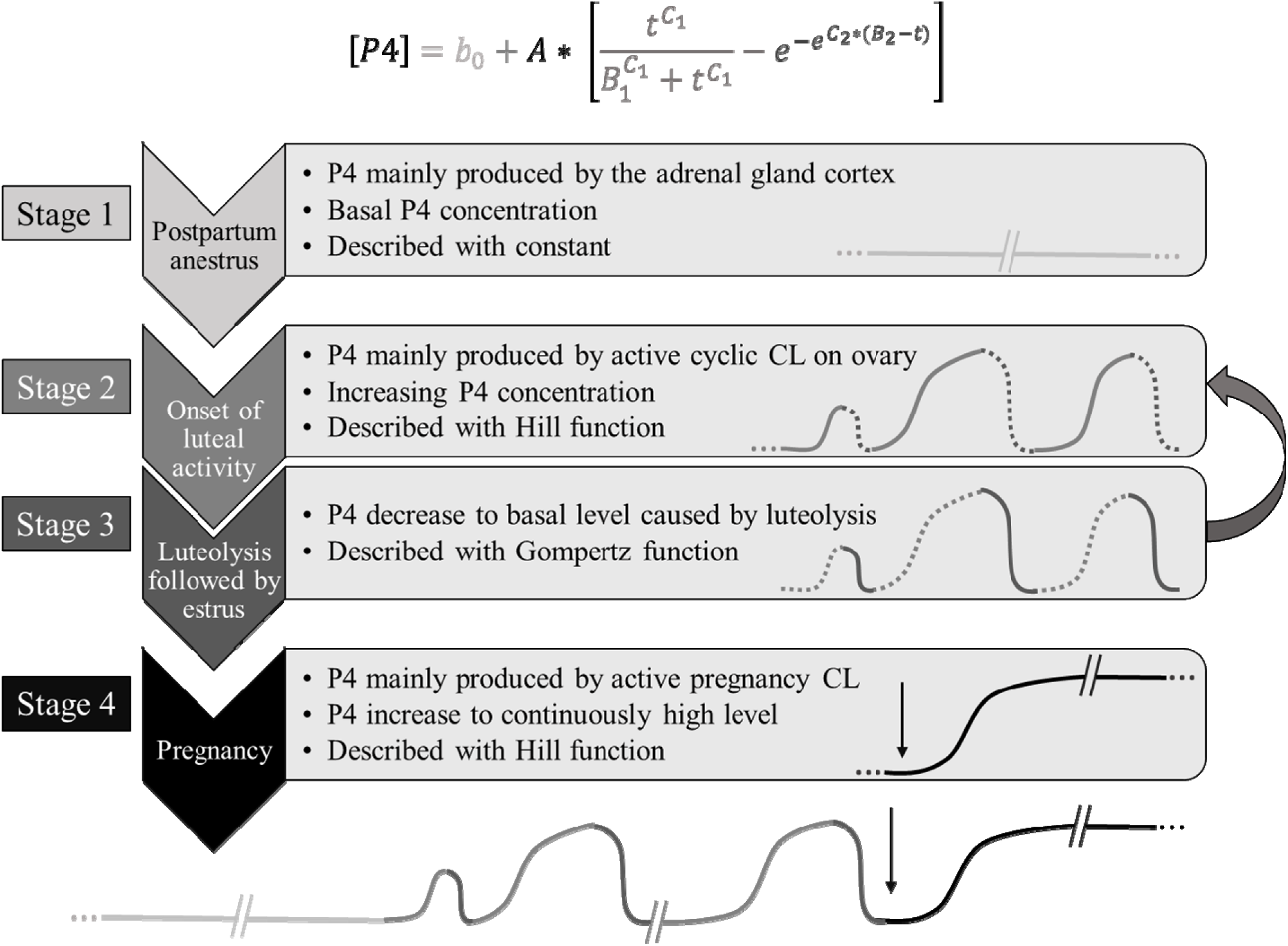
Different stages considered by the online monitoring system, i.e. *postpartum* anestrus (light grey, stage 1), cycling (grey, luteal phase, stage 2 and luteolysis, stage 3) and pregnancy (black, stage 4) and their link with progesterone (P4,) the physiological characteristics and the respective part of the mathematical model.

Several events can disrupt the expected course of P4 in the different stages. When the onset of cyclicity is delayed, *postpartum* anestrus will be prolonged, detected by a lag in the first luteal activity after calving. In the same way, the presence of follicular cysts or unexpected anestrus during the cyclic phase can be noted after luteolysis. The typical duration of the follicular phase in which no active CL produces P4 and accordingly, P4 is low, is 5-10 days (Royal et al., 2000). Longer periods of luteal inactivity after a detected luteolysis can indicate a problem for which a treatment may be required. A luteal cyst or a persistent CL will prolong the luteal phase of a cycle, because luteolysis is prevented. Contrarily, when an insemination did not result in gestation, the present CL will regress after 15 to 40 days depending on whether or not conception occurred, and a new attempt for pregnancy can be made.

### Passing the different stages

To determine the start of a new stage, an automated interpretation of the data is needed. The first stage ends with the onset of luteal activity corresponding to an increase in milk P4 is detected following a fixed rule, namely 3 measurements above 5 ng/mL within 3 days. A similar rule was used by Bruinjé et al., (2017b) who defined the onset of luteal activity as the first out of two successive measurements above 5 ng/mL following a measurement below 5 ng/mL. This rule is considered sufficiently accurate, because it is of no primary interest to know the exact starting moment of luteal activity on-farm. The end of stage 1 corresponds with the beginning of stage 2, being luteal activity.

In stage 2, the model is only allowed to describe the increase in P4 during luteal development. Onset of stage 3 (i.e. P4 decrease preceding estrus) of the abovementioned process requires reliable detection of luteolysis and is the key part of the developed system. Therefore, its principles are elucidated in Figure 2 where the different steps are presented. In general, we modified a ‘statistical control chart **(CC)** for individual subjects’ as described in Montgomery, (2013) to suit our system. Typically, the control limits of a CC for individual subjects are based on the moving range of successive measurements. However, we modified it to suit the need of detecting luteolysis, while being able to cope with the differing variability of the P4 measurements and possible outliers, and the large biological variation in the cycle characteristics. Overall, the CC consists of an upper **(UCL)** and lower **(LCL)** control limit, respectively given in eq. 2 and 3. These UCL and LCL are calculated initially on a training period as described below depending on the stage, and updated each time a new observation becomes available.

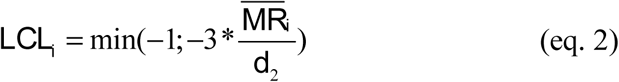

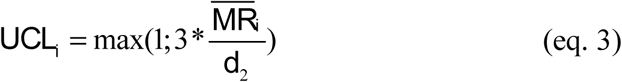

**Figure 2.**
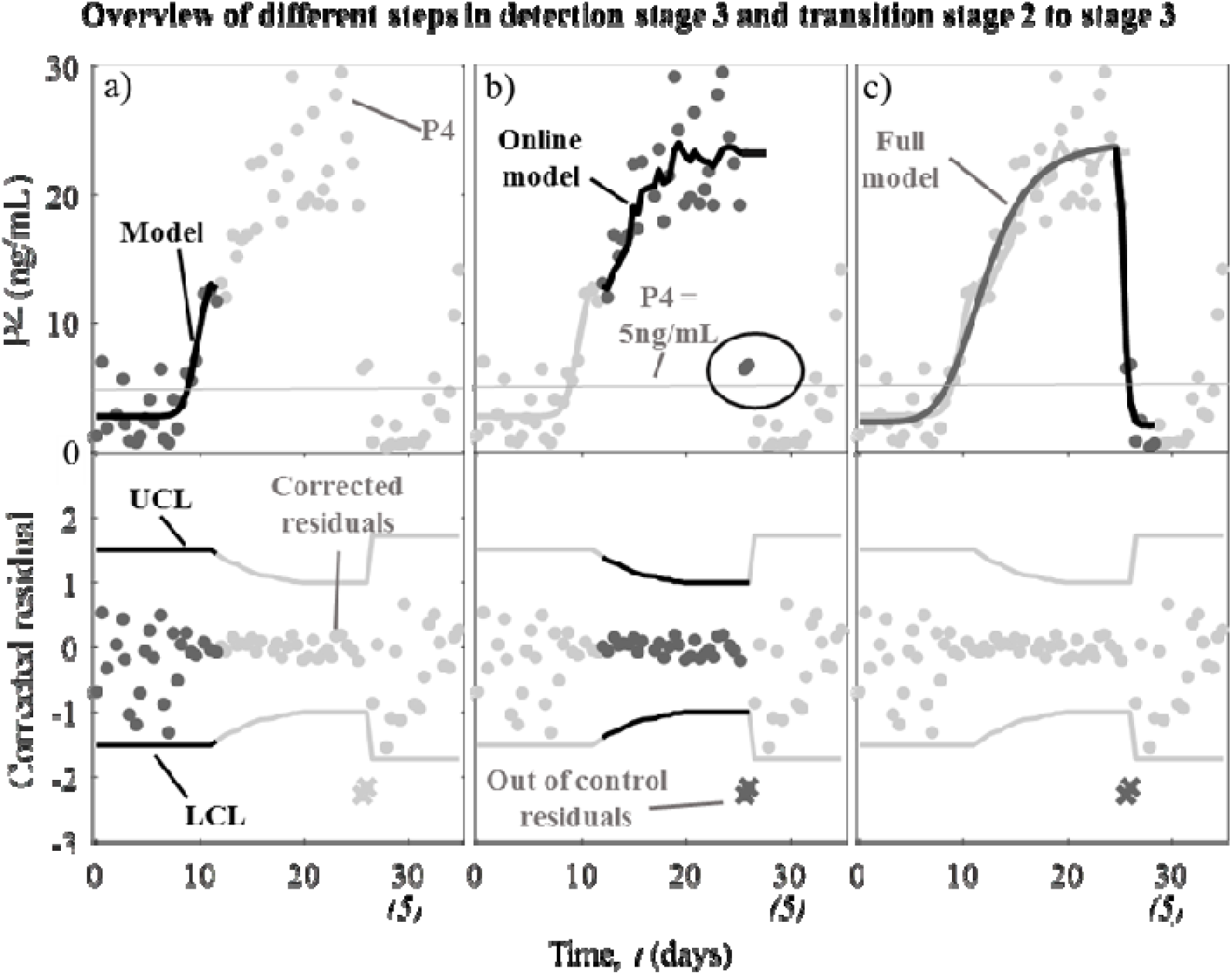
Overview of the online model and statistical control chart in the transition of stage 2 (onset luteal activity) to stage 3 (luteolysis). After a detected luteolysis (to), 10 days of data collection follow where after the increasing part of the model (stage 2) is fitted on the data, represented in panel a). The control chart for the current cycle consisting of the corrected residuals, an upper (UCL) and a lower control limit (LCL) is initiated. Each time a new measurement becomes available, the model parameters are updated and the corrected residual calculated accordingly (panel b). If a residual is out of the lower control limit, the model parameters are fixed to the previous update. At the third measurement out of control (x), stage 3 (luteolysis) is initiated and the decreasing Gompertz function added to the model (panel c). For this cow, the threshold of 5 ng/mL was not yet reached despite the confirmed luteolysis. The full model and its characteristics can now be calculated.

In eq. 2 and 3, the 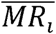 is equal to the mean moving range **(MR)** from the second observation until timepoint *i* of that cycle for each pair of observations, starting from *t*_0+1_ for each pair of observations *j* and *j*-1 given in eq. 4.

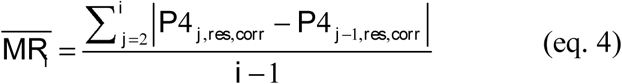

with *P4*_*j*,*reSiC*,*corr*_ as defined in eq. 5. Only values within the control limits of a cycle are included in the analysis. The *d*_*2*_ value for MR of paired observations is 1.128 (Montgomery, 2013). The minimum width of the control limits was set to 1 to avoid that phases with very low variability in the corrected residuals would make the CC too sensitive, e.g. due to saturation at very high P4 concentrations.

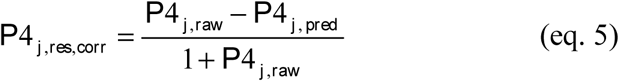

*P4*_*j*,*res*,*Corr*_ represents the corrected residual at time point *j*. *P4*_*i*,*raw*_ is the measured P4 and *P4*_*j*,*Pred*_ is the online predicted P4 at time point *j* by the Hill function describing stage 2. The correction factor is included to account for the unequal variance in the P4 data which is typically proportional with the P4 level (see also Friggens et al., 2008; Adriaens et al., 2017). The constant “1” was added to avoid the denominator being smaller than one, which would cause a large unbalance in the data. To make the concept more clear, the concrete steps are represented in Figure 2. These steps are repeated each time a new cycle is initiated by a stage 1 to stage 2 transition or a detected luteolysis (t0):

> Panel a). Directly after the transition from stage 1 to stage 2, or a detected luteolysis, 10 days of new data are collected to initiate the CC. The stage 2 model (i.e. increasing Hill function + baseline) is fitted on the available data, and the corrected residuals are calculated as described in eq.5. The calculated UCL and LCL are applied on the full 10-day initiation period and out-of-control measurements are evaluated. P4 data corresponding with residuals below the LCL are excluded from the calculations and the fixed initiation period control limits are updated if necessary.
>
> Panel b). After the initiation period, the model and control limits are updated each time a new P4 measurement, in-control according to the previous model parameters, becomes available. When luteolysis occurs, the corresponding measurements are consistently below the LCL and thus the model is not updated. When a third consecutive measurement is below the LCL, ‘luteolysis’ is considered confirmed at the first out-of-control measurement and stage 3 is initiated.
>
> Panel c). At the moment when the third consecutive measurement is below the LCL, the decreasing Gompertz function is added to the model, and the parameters of the full (Hill + Gompertz) model can be calculated using all data available for the current cycle. At this moment, a farmer can be informed that the cow is likely to become in estrus and might be inseminated after waiting a certain period. At this moment, the cycle restarts and the first step is repeated as shown in panel a).

Figure 3 shows for one example cow that the corrected residuals meet the preconditions needed for data to be used in a CC, namely homoscedasticity, normality, stationarity and the absence of significant autocorrelation. Especially, the homoscedasticity requirement (i.e. constant variance) is affected by the correction factor. In Figure 3, the raw residuals are shown as crosses and the corrected as circles. The upper panel a) shows the residuals reflected against the fitted full model representing the ‘average’ P4 level. When the P4 level is low, the raw residuals have a low variance and are close to zero. In contrast, when P4 is high, the raw residuals show a large range and more variability. The applied correction flattens out this difference without limiting the sensitivity for low P4 values, shown by the circles varying within a constant range around zero. Stationarity is proven by the fact that none of the time-series of online corrected residuals had an intercept nor slope significantly different from zero (average *p*-value = 0.404). The Anderson-Darling test pointed out that for none of the cows, the corrected residuals were data extracted from a normal distribution (average *p*-value < 0.001). Both the normal probability plot (b) and the histogram (c) show that small residuals are overrepresented with its nonlinear pattern and deviating Gauss-shape, respectively. Despite that normality is the least important condition for the CC to function properly (Montgomery, 2013), the overrepresentation of small residuals in our CC would lead to exceedingly narrow control limits. Therefore, the minimum width of 1 was set on the CC limits, as indicated in eq. 2 and 3. The last panel of Figure 3 shows the sample autocorrelation plot (d) for 20 lags and its corresponding boundaries (Mertens et al., 2009). The lack of systematic crossing of these boundaries, both within a profile and between profiles, indicates the absence of significant sample autocorrelation in the data.

**Figure 3.**
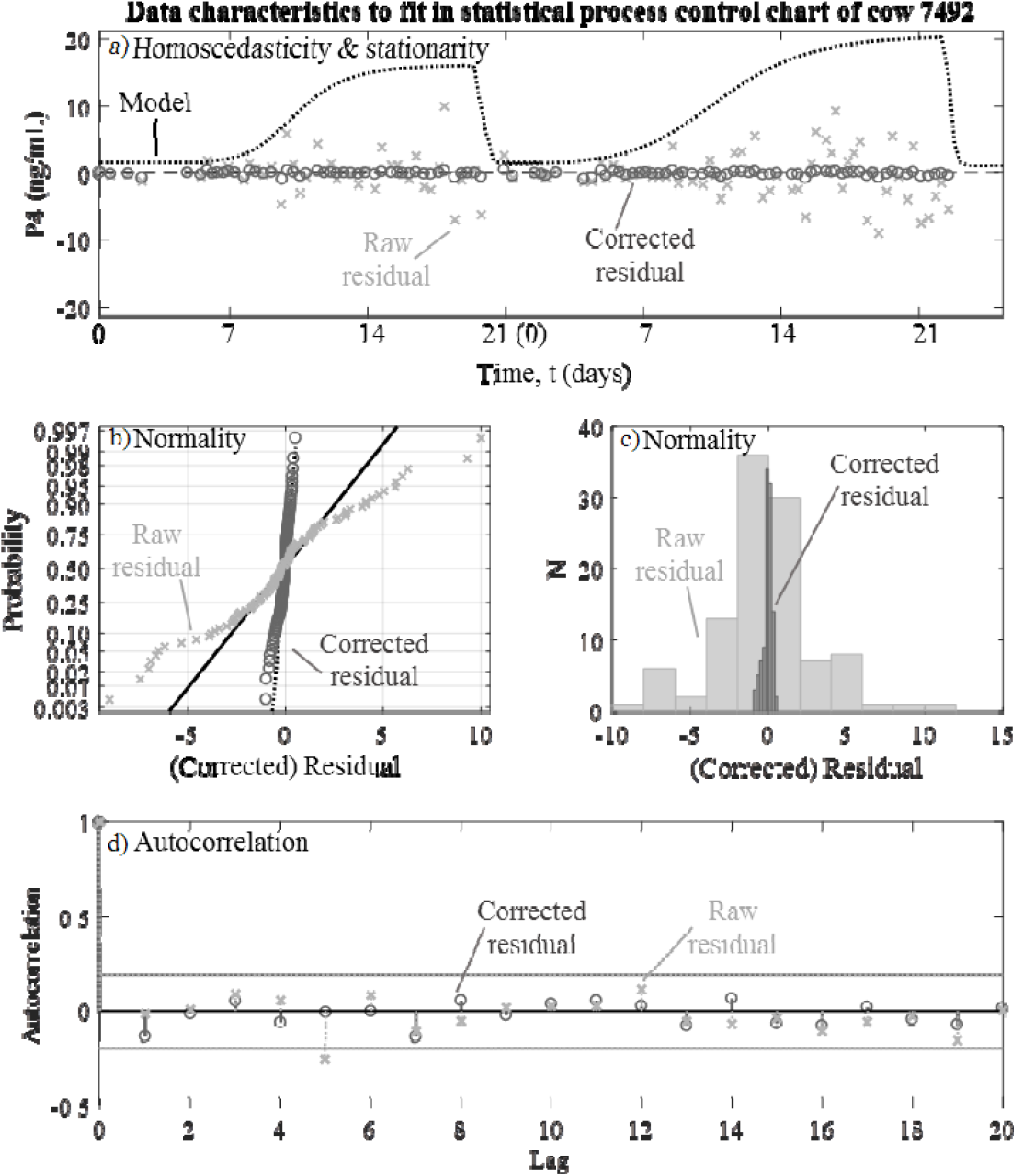
a) the uncorrected residuals of the online model show the heteroscedasticity of the P4 data proportional to the P4 level, represented by the model. The middle panels show the overrepresentation of small values in the corrected residuals, for which the minimum width of the control limits was set to 1. No significant autocorrelation, as shown in panel d is present in the data.

The onset of the last stage i.e. pregnancy, is characterized by the absence of luteolysis within a predefined window following insemination.

The combination of data-engineering (model step) and a statistical process control chart is known as ‘synergistic control’ (Mertens et al., 2009; Huybrechts et al., 2014). Therefore, our approach was named ‘**P**4-based **M**onitoring **A**lgorithm using **S**ynergistic **C**ontrol’ **(PMASC)**. All data processing was done in Matlab 2017a (Mathworks Inc., Natick, MA). The model was fitted and the parameters updated using a trust-region-reflective algorithm to obtain the least squares solution with a lower boundary of 0 for all the model parameters (cf. Coleman and Li, 1996).

### Monitoring fertility

The input of our online monitoring system is a time-series of P4 data of the same individual, further referred to as ‘P4 profile’. The CC-detected luteolyses were compared with the available reference data for onset of estrus, detected as described above. This means that a ‘confirmed estrus’ should have been preceded by a detected luteolysis, which on its turn was followed by a consistent period of low P4 during the follicular phase of the cycle. Additionally, the cows which were inseminated during this follicular phase and which did not experience a new luteolysis for more than 25 days after the previous one, were assumed pregnant. Pregnancy was confirmed by ultrasonography 40-50 days after insemination. Ideally, the cow would be assumed pregnant if there would be no luteolysis in a period of at least 35 days after insemination. However, the trial of 2016 ended earlier than 35 days after insemination for some of the cows. As we were not able to extend the data collection, this period was reduced to 25 days, potentially having a negative impact on the specificity for pregnancy detection. Additionally, when an abnormality was noted, the obtained profile characteristics were also studied to verify whether the system can manage abnormal P4 patterns.

## RESULTS AND DISCUSSION

### Monitoring fertility – normal profiles

An overview of the luteolyses detected by the system and either or not confirmed using method 1, 2, or both methods is shown in Table 2. In total, 50 luteolyses across 34 P4 profiles classified as ‘normal’ were detected by the system.

**Table 2.** Overview of the detected estruses and applied reference methodology to confirm them.

| # estruses detected by the algorithm and reference methods used in practice, normal profiles |  |  |  |  |  |  |
| --- | --- | --- | --- | --- | --- | --- |
| Year | # cows | M1<sup>1</sup> only | M2<sup>2</sup> only | M1+M2 | No reference | Total |
| 2016 | 15 | 6 | 1 | 6 | 5 | 18 |
| 2017 | 19 | 14 | 2 | 6 | 10 | 32 |
| **Total** | **34** | **20** | **3** | **12** | **15** | **50** |

<sup>1</sup> Method 1: uterus tonus, preovulatory follicle, CL; <sup>2</sup> Method 2: standing heat

For 15 of these detected luteolyses and subsequent follicular periods, we weren’t monitoring the cows to collect reference data, nor did the herdsmen note standing heat behavior. These cases thus occur in the first weeks of the P4 sampling periods when we didn’t have the cows synchronized yet (2016) or we weren’t monitoring the HN-P4-drop (2017). The other 35 cases were confirmed using method 1 or method 2 (see above). Briefly, for each case in which estrus was validated independently according to the reference methodology, the system provided timely detection of the luteolysis preceding ovulation. However, it should be noted that the number of cows for which there was reliable data available was rather limited. These 35 cases are not enough to draw strong conclusions on the usefulness of the system. However, within the data available, this is a very good result which needs to be confirmed through further evaluation of the PMASC on larger datasets.

For thirteen cows, a P4 decrease was not detected by the system within at least 25 days after insemination. Although 25 days might be too early to confirm pregnancy with high certainty, the absence of such a decrease confirms absence of a new estrus. It was shown before that the first 32-35 days of a pregnancy are crucial (Chebel et al., 2004; López-Gatius, 2012; Ricci et al., 2017), and accordingly, it is recommended to monitor the P4 profile for at least this period, and preferably longer. Twelve of these cows were confirmed to be pregnant in week 6 of gestation. One cow turned out not to be pregnant at that time, but there was no further P4 data available to confirm early embryonic death or a prolonged luteal phase as the milk P4 monitoring period stopped 26 days after insemination. The first estrus detected visually by the herdsmen for this cow was 59 days after the previous one noted by the PMASC system. Accordingly, this at least indicates that the developed system can deal with prolonged luteal phases after insemination, which could be related to pregnancy.

Next to identifying luteolysis and pregnancy, the model can also be used to calculate cycle characteristics from the model parameters. As such, the baseline and luteal P4 levels, slopes and lengths of the different phases can provide monitoring information. For example, the average baseline concentration (*b0*) of the P4 profiles in this study was 1.9±0.7 ng/mL (mean±SD), while the average maximal P4 concentration in the luteal phase (*b0*+*A*) was 20.9±6.1 ng/mL, which is in accordance with the profile characteristics reported by Blavy et al. (2016). The average slope of the increasing part of each cycle (3.1±2.5 ng/mL per day) is less steep compared to the decreasing part (31.2±31.8, ng/mL per day). As such, the time in which the P4 level increases from a mean baseline of 1.9 ng/mL to a mean maximal level of 20.9 ng/mL is on average 6.05 days. In contrast, the time in which the P4 level drops from maximal to basal P4 concentrations is only 15.5 hours on average, indicating the high rate at which the P4 levels drop after luteolysis. Both observations are in line with previous research on plasma P4 concentrations (Mann, 2009; Meier et al., 2009).

Although good results were obtained with the PMASC, the following aspects of the methodology might need further investigation and optimization: (1) the apparently arbitrary use of the rule to start stage 2; (2) the need for 3 observations out of control for confirmation of luteolysis and (3) the use of a complete, once-a-milking sampling scheme. The first aspect concerns the transition from stage 1 (postpartum anestrus) to stage 2 right at the onset of first luteal activity. Since our methodology does not apply smoothing, raw P4 measurements determine the start of luteal activity and accordingly, the increasing part of the model. As a result, when the raw data accidentally fulfils the conditions set for initiating stage 2, the increasing model might be fitted and updated while there is no actual luteal activity yet. Nevertheless, in that case, the residuals of the model are typically not large enough to provoke an indication of luteolysis in the CC. Consequently, this is not causing any problems in the studied cycles. When after a certain period luteal activity properly starts, the model parameters will adapt and the correct information can be extracted. Additionally, it should be noted that the first ovulation of a cow has already passed at the time the system detects the first luteal activity after calving. This is a general drawback of using P4 for fertility monitoring, because luteolysis does not precede first ovulation and therefore the onset of follicular activity cannot be detected. Nevertheless, common dairy practice advises to not inseminate on this first ovulation after calving, thereby mitigating this drawback (Peter et al., 2009).

The second aspect concerns the use of 3 measurements below the LCL to confirm luteolysis and inform the farmer about a cow that comes in heat. Timely informing a farmer is necessary to enable optimal insemination timing, 24 to 12 hours before ovulation (Roelofs, 2005). However, Roelofs et al. (2006) also showed in their study that milk P4 dropped below a 5 ng threshold at least 54 hours before ovulation. Accordingly, we decided that using 3 measurements to assure luteolysis is still in time, thereby minimizing the risk of false alarms. Concrete, we took into account that the time between milkings is generally 14 hours or less. This means that luteolysis is notified by the algorithm within 28 hours after the actual drop in P4 and on average about 30 hours before ovulation (Roelofs et al., 2006), which is well in time. As such, the corresponding time lag on the notification of luteolysis does not outweigh the advantage of certainty, and does not hamper the timely indication of the optimal insemination moment. Moreover, the indication of the moment of luteolysis itself is accurate, being independent of the time lag. Both luteolysis, which is at the moment of the first of 3 consecutive measurements with a residual below the LCL, as well as the model characteristics can be used to predict the moment of ovulation. The latter possibly provides a large advantage compared to the use of a fixed threshold on a smoothed P4 profile, for which some difficulties are identified. For example, the smoothing introduces an unpredictable time lag on the moment at which the smoothed P4 profile drops below the fixed threshold, influenced by the sampling frequency and the rate of P4 decrease (Friggens et al., 2008; Bruinjé et al., 2017b). This might not influence detection of the follicular phase itself, but complicates the proper prediction of ovulation based on the moment of luteolysis. Using direct mathematical modeling of the P4 cycles, luteolysis might be characterized in a more consistent way. For example, in Figure 2 b) the 2 measurements identified by the PMASC as out of control do not undercut a fixed threshold of 5 ng/mL. Still, the luteolysis is detected as soon as the P4 concentration drops, independent of any predetermined fixed decision rules. Additionally, Friggens and colleagues (2008) observed that sometimes a higher threshold (e.g. 6 ng/mL) is needed to identify ‘high P4 estruses’. This observation supports the idea of using an individualized approach rather than fixed thresholds. Accordingly, future research should further investigate the relation between luteal decay and ovulation and factors affecting this.

This brings us to the third aspect of the discussion on the use of maximum sampling frequency. In this study, all available data i.e. P4 measured 2-3 times a day were included in the analysis. However, to obtain a cost-effective system, adjusting the sampling frequency dependent on the progress of the cycle would be preferred, as already implemented in the currently used online monitoring algorithm (Friggens and Chagunda, 2005; Friggens et al., 2008). The principles from their approach might easily be integrated in the PMASC. For example, in the increasing phase of each cycle, which typically lasts for 10 – 20 days, the sampling frequency can be reduced to once per two days or even less. When luteolysis is expected or when 1 measurement undercuts the LCL, the sampling frequency could be increased to once a day or at each milking. Future research should focus on the consistency of the model parameters and characteristics when sampling frequency is adjusted, but it is assumed that this system might be more robust against missing values in its decision making, compared to a smoothed level and fixed threshold. Depending on the practical decisions in this context and on-farm implementations, the PMASC monitoring system should be optimized accordingly.

### Monitoring fertility - abnormal profiles

It is important to illustrate that the developed PMASC has no problems with deviating fertility courses, which would limit the use of the monitoring system to model-profiles only. Therefore, a few cases of fertility issues and the corresponding models are discussed in this paragraph. It is by no means our intention to describe at this point how the system can detect these fertility issues, because the number of aberrant cases was too limited for a thorough validation. The conceptual framework described in Friggens and Chagunda, (2005), in which the risk of luteal and follicular cysts is calculated based on the length of the luteal and follicular phases, could be applied on the profile characteristics calculated from the proposed approach as well. For example, the length of the follicular phase is typically 5 to 10 days. When the P4 model does not reach luteal concentrations within an acceptable time period based on this knowledge, the risk of follicular cysts increases. The same reasoning can be applied to the luteal phase for luteal cysts or persistent CL.

To illustrate this concept and the applicability of our system for the detection of aberrant profiles, a P4 profile and the respective monitoring of a cow suffering with cystic ovary disease during the trial is shown in Figure 4. After detection of the follicular cyst, the cow was immediately treated with a GnRH-agonist inducing ovulation on day 22. Prior to the GnRH treatment, P4 was low and the corresponding baseline concentration varied around 1.5 ng/mL for more than 20 days. No information was available for the preceding period, but based on these data, the follicular phase lasted for at least 22 days. At that time, this cow had already been 103 days in lactation and seen in heat by the herdsmen 3 times. From the treatment day on, the milk P4 concentration increased slowly towards an intermediate luteal concentration of 12 ng/mL. As this was a cow flagged for close monitoring, she was scanned repeatedly, and in the morning of the 27^th^ day after GnRH treatment, the presence of a luteal cyst was confirmed. At this moment, the cow was treated with *Dinoprost tromethamine*, triggering an immediate luteolytic reaction, reflected in luteolysis detected by the system in the afternoon on that same day. Although the P4 concentration of this cow was rather unstable, reflected in the high variability in the measurements during the luteal cystic phase, the system was able to identify the correct overall P4 pattern and as such, might be used to inform the farmer on deviating profiles and corresponding problems.

**Figure 4.**
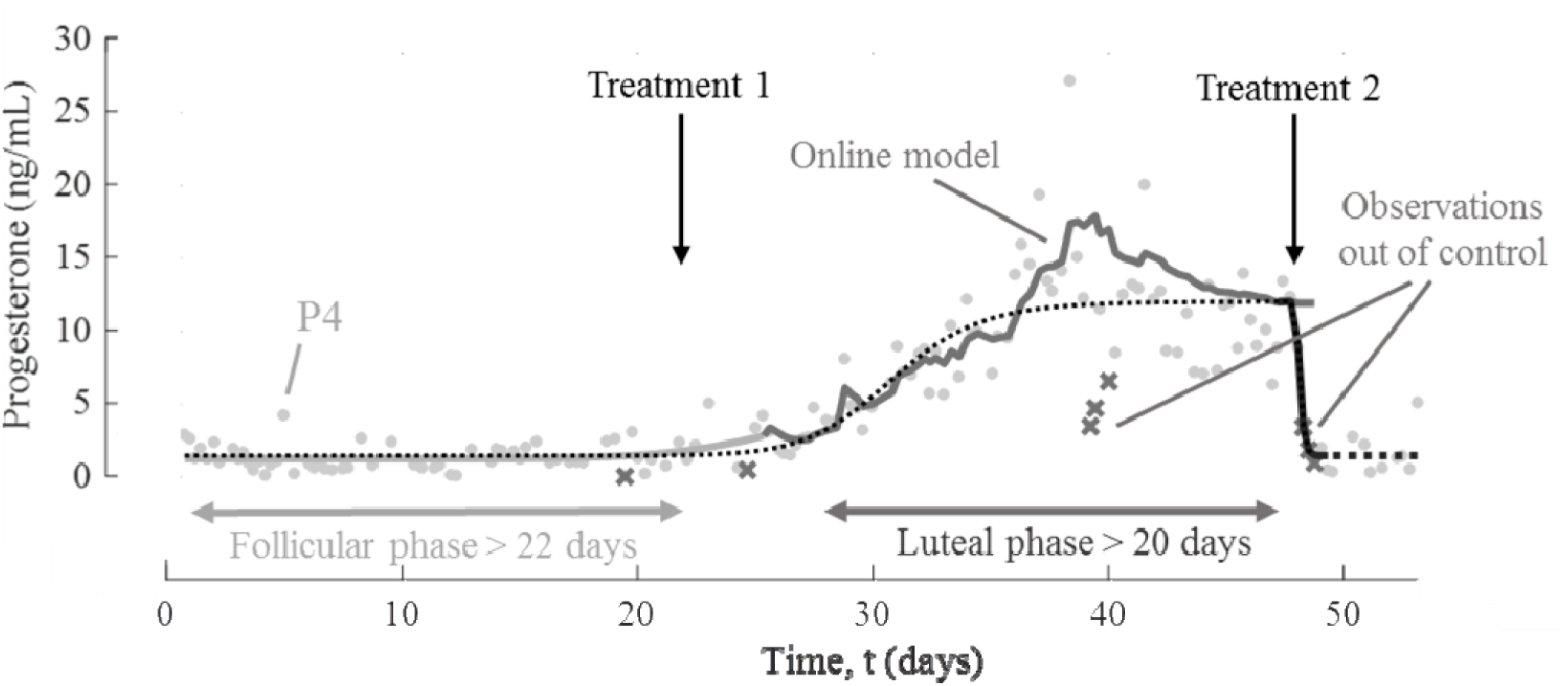
Example of the P4 profile of a cow with cystic ovary disease. At the start, this cow developed a follicular cyst after a normal cycle. This was confirmed on t_22_ and she was treated with a GnRH-agonist. Hereafter, the cow developed a *corpus luteum* (CL) producing P4, which evolved to a luteal cyst. This was treated with *Dinoprostum tromethamini* on day 49, immediately followed by luteolysis detected by the system. During her luteal cystic period, the P4 production seemed rather low and unstable, reflected in the residuals crossing the LCL boundary e.g. around day 40. As these are interrupted with a high measurement, no luteolysis attention is triggered.

In total, from the 5 follicular cysts detected during the trials, 3 were preceded by luteolysis and indicated by the monitoring system. For the other 2, the cyst was already present at the start of the trial and the proceeding luteolysis was outside the range of the dataset. Two luteal cysts were detected from which one evolved to a follicular cyst spontaneously (serum-P4 went from 4.6 ng/mL to 0.2 ng/mL). An overview of these cases is given in Table 3.

**Table 3.** Overview of the detected abnormalities

| Year | # cows | Follicular cyst | Luteal cyst | Anestrus |
| --- | --- | --- | --- | --- |
| 2016 | 1 | 1 | 1 | 0 |
| 2017 | 5 | 4 | 1 | 1 |
| **Total** | 6 | 5 | 2 | 1 |

Although only a few cases were available and thus precaution should be taken when drawing conclusions, it was noted that the maximal luteal P4 levels for the cows suffering from fertility issues estimated by the model was significantly lower compared to the normal profiles (15.2 ng/mL vs. 20.9 ng/mL respectively, *p*-value = 0.0197). This is in accordance with the observation of Hatler et al. (2003) and Rosenberg (2010), who reported that cows diagnosed with ovarian follicular cysts had intermediate concentrations of P4. This can be caused by an inferior quality of the CL. As seen in Figure 4, an irregular pattern of the milk P4 is noted. Several measurements undercut the LCL in the luteal phase of the cycle. Because these are interrupted by in-control measurement, they do not trigger the alarm for luteolysis. It seems likely that our system reacts more quickly and sensitively on actual P4 concentration changes compared to smoothing techniques. However, because most fertility problems affect the P4 profile on a long term, the real effect on detection rates should be further investigated by involving more cases and exploring the differences in profile characteristics and their predictive value for abnormal profiles.

## CONCLUSIONS

We developed an innovative system for dairy cow fertility monitoring based on milk P4. This system is an alternative for the currently used filtering techniques and fixed thresholds for estrus detection, and its associated problems. It consists of a mathematical model describing the P4 profile combined with a statistical control chart for the detection of luteolysis. The model characteristics and the control chart have the potential to identify first luteal activity, estrus, pregnancy, early embryonic death and abnormal profiles indicating ovarian problems. As the approach is rather straightforward, it allows the implementation in an online monitoring system for on-farm use. Additional research is required to optimize sampling frequency, clarify the link between the model characteristics and the occurrence of abnormalities and to investigate the exact relation between the P4 profile and the time of ovulation.

## ACKNOWLEDGEMENTS

This work was supported by the Institute for the Promotion of Innovation through Science and Technology in Flanders, Belgium (IWT) [IWT-LA project 110770]. Ines Adriaens and Ben Aernouts are supported by the Fund for Scientific Research (FWO) Flanders, respectively grant number 11ZG916N and 12K3916N. We thank Carmen Adriaens for critical reading of the manuscript.

